# The human oral phageome is highly diverse and rich in jumbo phages

**DOI:** 10.1101/2020.07.06.186817

**Authors:** Victoria R. Carr, Andrey Shkoporov, David Gomez-Cabrero, Peter Mullany, Colin Hill, David L. Moyes

## Abstract

Until recently, the contribution of bacteriophages to the composition and function of the human microbiome has been largely overlooked. Recent developments in discovering novel bacteriophages from human metagenomes have been mostly focused on the gut. Here we profile and compare the phageome of 633 human oral sites and 221 paired gut phageomes acquired from countries worldwide. Phageome profiles are specific to oral sites and the gut across individuals, with the largest separation between the oral cavity and the gut. The differences in phage genotypes is bolstered by distinct phage host compositions between these sites. The greatest diversity of phages is found on the tongue and in dental plaque. Notably, 37 unique circular jumbo phage genomes were identified in oral sites, particularly on the dorsum of the tongue, whilst none were found in the gut. Oral sites provide conducive environments, such as robust biofilms, that can harbour genetically diverse phages.

## Introduction

Bacteriophages are the most abundant viral components of the human microbiome. They are viruses that infect and replicate their genome within bacterial or archaeal cells, and are likely to have significant effects on microbial composition and function^1,2^. Like eukaryotic viruses, they can have single or double stranded DNA or RNA genomes. They also have two principal life cycles: virulent, which destroy bacterial cells immediately after replication; and temperate, which have the option of integrating their genome into the host genome (lysogeny) and are involved in the horizontal gene transfer of many genetic elements, including virulence factors.

Many studies that attempt to profile bacteriophages in human microbiomes use computational analysis of faecal metagenomic data, often following enrichment of virus-like particles^3,4^. This has led to the discovery of hundreds of thousands of novel and uncultivated virus genomes^5^. These include crAssphages, a highly abundant bacteriophage clade currently thought to play a special role in human and primate gut microbiomes^6–8^; the discovery of atypically large bacteriophages greater than 200 kb in length, known as jumbo phages^9^; and more recently megaphages with genomes larger than 500 kb^10^. Only a handful of studies have profiled bacteriophages using metagenomic data from the oral cavity^11,12^. The heterogeneity of bacteriophage genomes, and lack of correlation between phage phylogeny and that of their hosts, makes classification and host assignment challenging tasks, leaving a relatively unexplored melting pot of “viral dark matter” ^3,13^.

Here, we profile bacteriophage DNA from different locations in the human gastrointestinal tract, specifically comparing gut (represented by faecal samples) with paired saliva and dental plaque from China, and the dorsum of tongue, dental plaque and buccal mucosa from longitudinal samples from the USA. We identify novel bacteriophages, including jumbo phages, from assembled metagenomic contigs using *de novo* bioinformatic pipelines, including viral motif recognition^14^ and protein-coding gene-sharing networks^15^ to identify and classify linear and circular viral contigs. Bacteriophage hosts are also predicted using CRISPR spacer matches with reference bacterial genomes^16^.

We report that bacteriophages and their host profiles are specific to gastrointestinal tract (GIT) sites, with more differences found between the oral cavity and the gut than between different oral sites. The dorsum of the tongue contains a greater diversity of bacteriophages than paired stool samples, saliva and buccal mucosa. Consistent with greater genotypic diversity, 37 unique circular jumbo phage genomes were found in oral cavities from China and the USA, mainly on the dorsum of the tongue from the USA, while none were identified in paired stool samples. In addition, bacteriophages can persist in some individuals for months at a time, including phage families that are common to both oral cavity and gut, or are specific to a GIT site, such as the crAss-like family of the gut, *Inoviridae* of the oral cavity, and *Microviridae* of the gut, dorsum of the tongue and saliva.

## Results

### Phage composition and diversity differs between GIT sites

A catalogue of 139,929 phage contigs were identified from 854 oral and gut metagenomes. Phage contigs were grouped into clusters using vConTACT2^15^, a software based on gene-sharing profiles that has been applied to stratify uncultured viral populations in a variety of environments, including human gut metagenomes^3,16^. 75,680 phage contigs were clustered into 14,078 phage clusters, while the remaining 64,249 remain as singletons. It is more common to find phage contigs that are unique to one sample than phage clusters found, meaning phage contig profiles at strain level are more variable than phage cluster profiles (Supplementary Fig. 1). β-diversity was evaluated with Bray-Curtis dissimilarity metric from phage cluster abundance profiles to investigate differences between metagenomes. Samples were clustered using unsupervised Silhouette analysis on k-medoids of non-metric multidimensional scaling (NMDS) dimensions. Four groups were generated (Fig. 1a). Most phage clusters are exclusively found in either Group 2, 3 or 4 or are shared between Group 2 and 3 (Fig. 1b). The composition of these groups is dominated by samples from a particular GIT site (Fig. 1c). Groups 1 to 3 contain all oral sites: Group 1 consists of mostly buccal mucosa, Group 2 contains mostly dorsum of tongue and saliva, and Group 3 has mostly dental samples. In contrast, Group 4 contains exclusively stool samples. These results reveal that many phage clusters are solely found in either dorsum of tongue (Group 2), dental (Group 3) or stool samples (Group 4), but most phage clusters that are shared occur in both dorsum of tongue and dental sites. Hierarchical clustering of the relative abundance of phage clusters also shows GIT sites can be differentiated by phage cluster profiles, especially between dental, dorsum of tongue and stool samples (Supplementary Fig. 2). GIT sites have distinct phage cluster profiles, and phage clusters that are shared are mostly found in oral sites. Incomplete metadata on sex and age meant that individual variation could not be considered (Supplementary Fig. 3).

**Figure 1:**
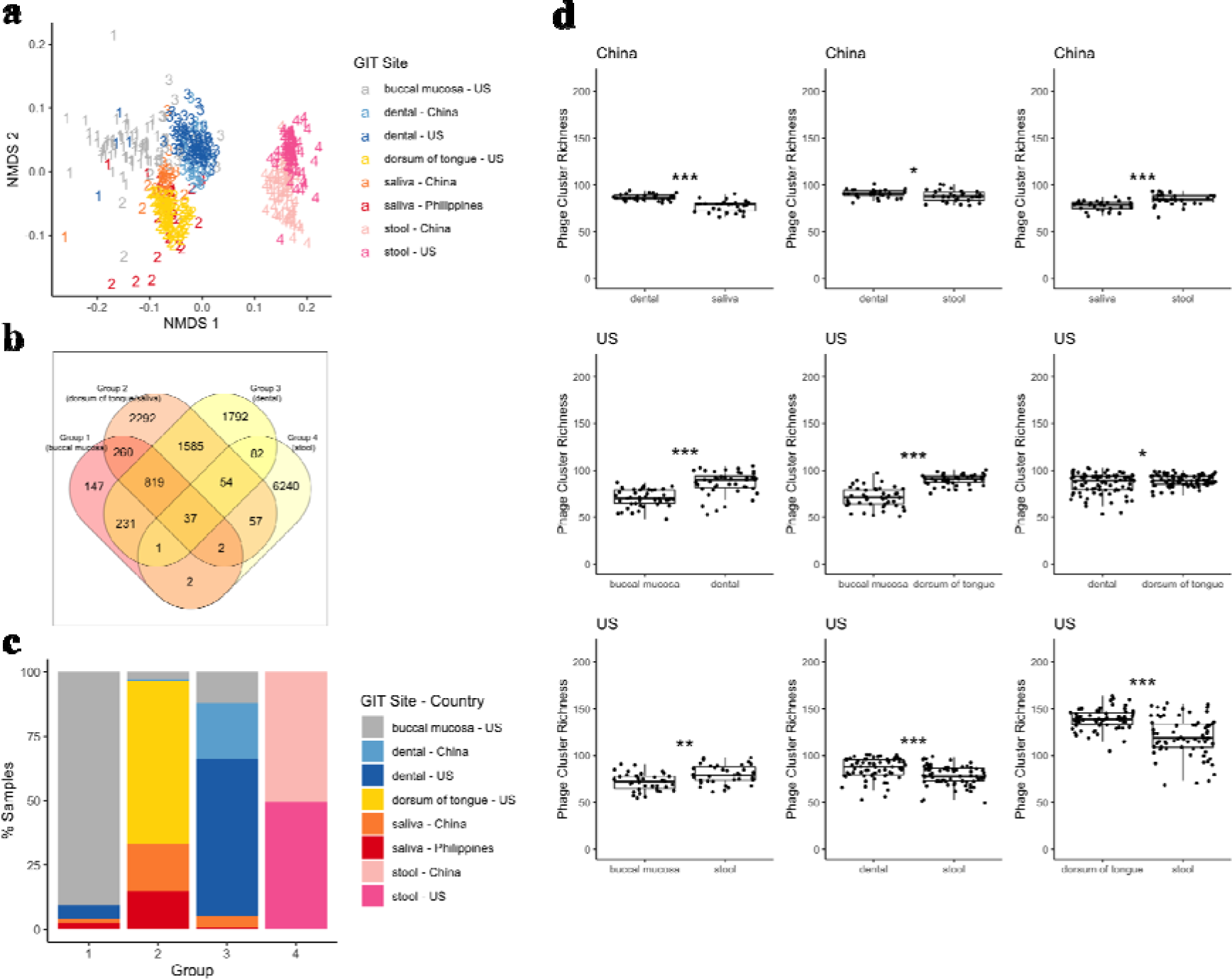
Phage incidence and abundance profiles. **a)** Non-metric multidimensional scaling of Bray-Curtis dissimilarity between phage incidence profiles of samples (excluding longitudinal USA). Ordination coordinates are grouped by k-medoids clustering, where number of groups, *k*, has the largest Silhouette width. USA buccal mucosa (n = 87), dorsum of tongue (n = 90), dental plaque (n = 90) and stool (n = 70); China dental plaque (n = 32), saliva (n = 33) and stool (n = 72); and Philippines saliva (n = 24). **b)** Number of viral clusters in each group. **c)** Percentage of samples in each group from 1a, labelled by GIT site and country. **d)** Phage Cluster Richness between paired GIT sites. Phage Cluster Richness is defined as the number of unique viral clusters in a sample that is subsampled to the smallest number of non-unique clusters. Phage Cluster Richness is calculated for samples of individuals from China (dental plaque and saliva: n = 30, stool and saliva: n = 30, stool and dental plaque: n = 30) and the USA (buccal mucosa and dental plaque: n = 45, buccal mucosa and dorsum of tongue: n = 45, buccal mucosa and stool: n = 36, dental plaque and dorsum of tongue: n = 86, dental plaque and stool: n = 67, dorsum of tongue and stool: n = 68) (excluding longitudinal USA) with Two-sided Wilcoxon Rank Sum Test (p < 0.05 as *, < 0.01 as **, < 0.005 as ***). Centre line is median, box limits are upper and lower quartiles, whiskers are 1,5x interquartile ranges and points beyond whiskers are outliers.

Additionally, the α-diversity of the phage cluster composition was compared between paired GIT sites from China and the USA. The Phage Cluster Richness, i.e. the number of unique phage clusters, was evaluated for every paired sample. Dental plaque from the USA (p = 2.84×10^−4^) and from China (p = 0.0476), and dorsum of the tongue samples from the USA (p = 1.89×10^−7^) have a significantly greater Phage Cluster Richness than stool samples (Two-sided Wilcoxon Rank Sum Test; Fig. 1d). USA dorsum of the tongue samples also have a significantly higher richness than dental plaque (p = 0.0469). Buccal mucosa from the USA and saliva from China have the lowest richness compared to all other paired GIT sites (saliva vs. dental plaque from China: p = 2.94×10^−6^, saliva vs. stool from China: p = 1.93×10^−4^, buccal mucosa vs. dental plaque from the USA: p = 3.91×10^−7^, buccal mucosa vs. dorsum of the tongue from the USA: p = 1.26×10^−8^, and buccal mucosa vs. stool from the USA: p = 0.00175).

Most oral sites and stool metagenomes contain a high abundance of *Siphoviridae, Myoviridae* and *Podoviridae* phage families but there are some distinctive differences in the less abundant phage families between GIT sites (Supplementary Fig. 4). Bacterial hosts for phages were predicted using Demovir (https://github.com/feargalr/Demovir). Although host bacteria could only be predicted for 11.8 % (16,513/139,929) of phage contigs, these highly abundant phage families infect a range of bacterial genera (Supplementary Fig. 5). crAss-like phage clusters are only found in stool samples and in *Prevotella* spp. as host, apart from two saliva and two dorsum of the tongue samples (Supplementary Fig. 6). In contrast to the crAss-like family, *Inoviridae* are found almost exclusively in oral sites and predicted to infect *Neisseria* species. *Microviridae* are present in dorsum of the tongue, saliva and stool samples but are far less prevalent in buccal mucosa and dental plaque. *Prevotella* in oral sites and *Faecalibacterium* are the only genera predicted to contain *Microviridae. Bicaudaviridae* are only found in ten dorsum of the tongue samples and are represented by only one phage cluster. There are 30 *Inoviridae*, 56 *Microviridae*, 6967 *Siphoviridae*, 3433 *Myoviridae*, 728 *Podoviridae* and 64 crAss-like phage clusters (Supplementary Fig. 6). Across highly abundant families (i.e. *Siphoviridae, Myoviridae* and *Podoviridae*), GIT sites are clustered by the incidence of phage clusters suggesting that phage clusters of the same family are also specific to GIT site. In terms of lower abundance families found across multiple GIT sites, there is a pronounced separation of oral sites and stool in *Microviridae* phage clusters, with some separation of oral sites by *Inoviridae* phage clusters. Notably, the *Microviridae* family is dominated by a few phage clusters in oral sites in contrast to stool that contains many different phage clusters.

### Phage hosts match varied microbial composition across GIT sites

Bacteriophages have previously been shown to modulate the microbiota composition (in particular the bacterial and archaeal composition) in the mouse gut via phage infection and lysis of specific host bacteria^2^. To investigate whether microbial composition is associated with distinctions in phageome profiles between GIT sites, bacterial and archaeal taxonomies were profiled using MetaPhlAn2^17^. Clustering of taxonomic composition is associated with GIT site, with greater separation between oral sites and gut (Fig. 2a). Procrustes analysis was applied to match corresponding points between phageome and taxonomic profiles. Procrustean randomisation test (PROTEST) was then used to determine whether two profiles showed significant association. Microbial composition and phageome profiles correlate and co-locate by GIT site, especially between stool and dental samples (Fig. 2b) (0.70 to a significance of p = 0.001 in PROTEST). Predicted phage hosts group by GIT site with very little overlap between gut and oral sites (Fig 2c, Supplementary Fig. 7). *Actinomyces, Atopobium, Campylobacter, Fusobacterium, Neisseria* and *Rothia* genera, that are mostly found in the oral cavity (Supplementary Fig. 8), are predicted hosts of oral site phage (Fig 2c). Likewise, *Bacteroides, Bifidobacterium, Clostridium, Roseburia, Ruminococcus* and *Parabacteroides* genera found more exclusively in stool samples are also potential phage hosts in the gut. However, *Eubacterium*, a bacterium mostly found in stool, and *Haemophilus, Prevotella, Streptococcus* and *Veillonella*, bacteria mainly residing in the oral sites, are prevalent across all oral sites and the gut. Upon closer inspection of the abundance of these genera at a species level, there are some species that are more prevalent in either the oral cavity or the gut. For instance, *E. brachy* and *E. saphenum* are species of *Eubacterium* that are found almost exclusively in the oral cavity (Supplementary Fig. 9). Likewise, *P. copri* mostly represents *Prevotella*, and is found almost exclusively in the gut. Generally, phage host predictions match microbial composition at a genus level.

**Figure 2:**
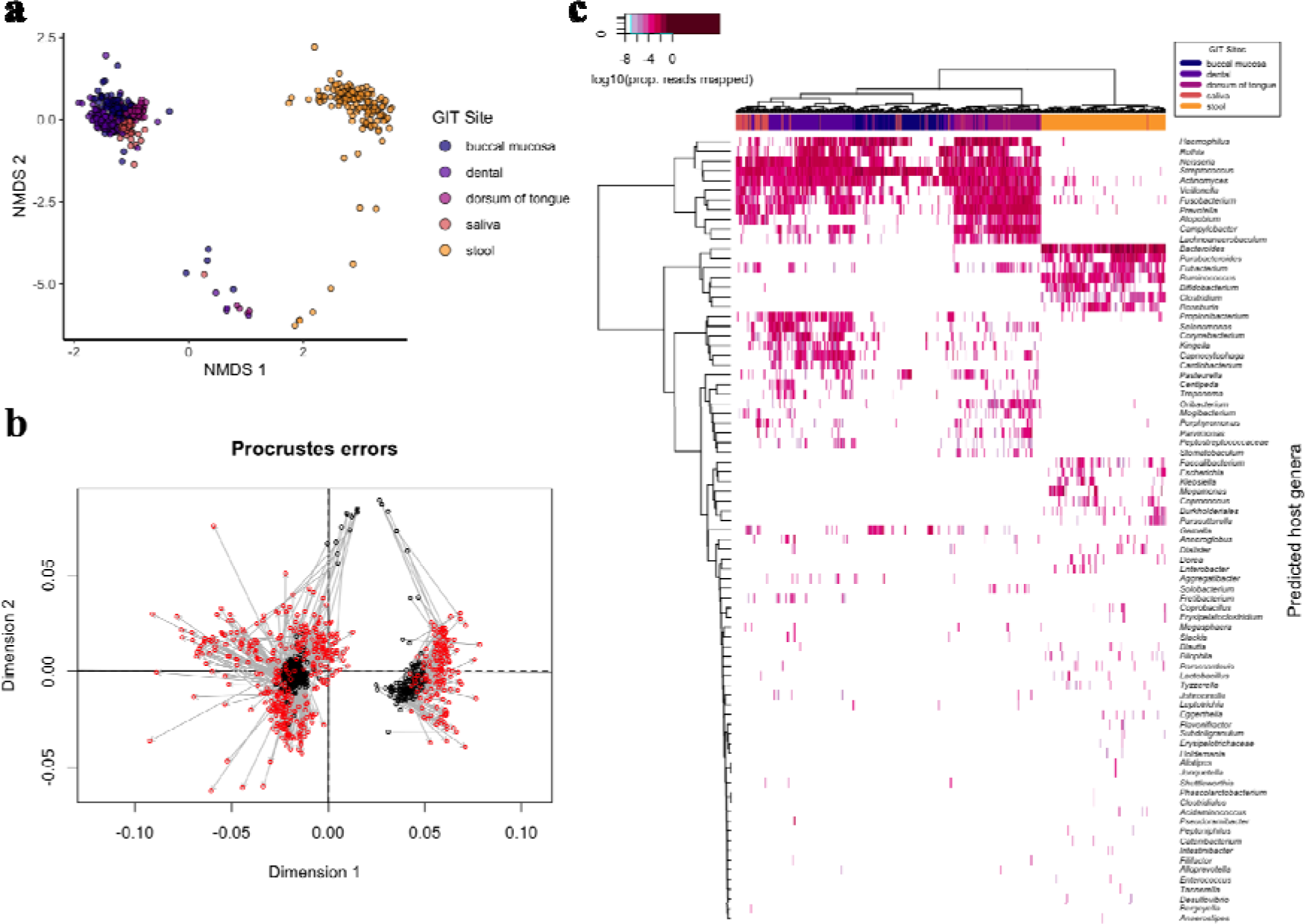
Relationship between phage profiles and microbial composition, and abundance of predicted phage hosts. **a)** Non-metric multidimensional scaling (NMDS) of Bray-Curtis dissimilarity between microbial taxa incidence of samples (excluding longitudinal USA and n = 21 samples with suspected contamination) labelled by GIT site. **b)** Procrustes rotation of NMDS coordinates between microbial genera profile from 2a (black) and phageome profile from 1a (red). Correlation in symmetric Procrustes rotation = 0.70 (p = 0.001; 999 permutations; PROTEST). **c)** Log10 of the proportion of reads mapped to phage contigs with predicted host for each sample, clustered by hierarchical clustering, x-axis coloured by GIT site and y-axis labelled by genus of predicted host. USA buccal mucosa (n = 87), dorsum of tongue (n = 90), dental plaque (n = 90) and stool (n = 70); China dental plaque (n = 32), saliva (n = 33) and stool (n = 72); and Philippines saliva (n = 24).

### Stability of phage clusters across longitudinal metagenomes

To clarify whether phages that are associated with a GIT site are also stable over time, phage clusters were profiled in longitudinal samples from the USA taken over a two-year period with a minimum of two and maximum of six sampling timepoints. The proportions of total phage clusters (Fig. 3a) and total reads mapped to these phage clusters (Fig. 3b) drops over time across all GIT sites, but this is unclear from timepoints 4-6 in buccal mucosa, 4 in dorsum of tongue and 5-6 in stool due to a small number of samples.

**Figure 3:**
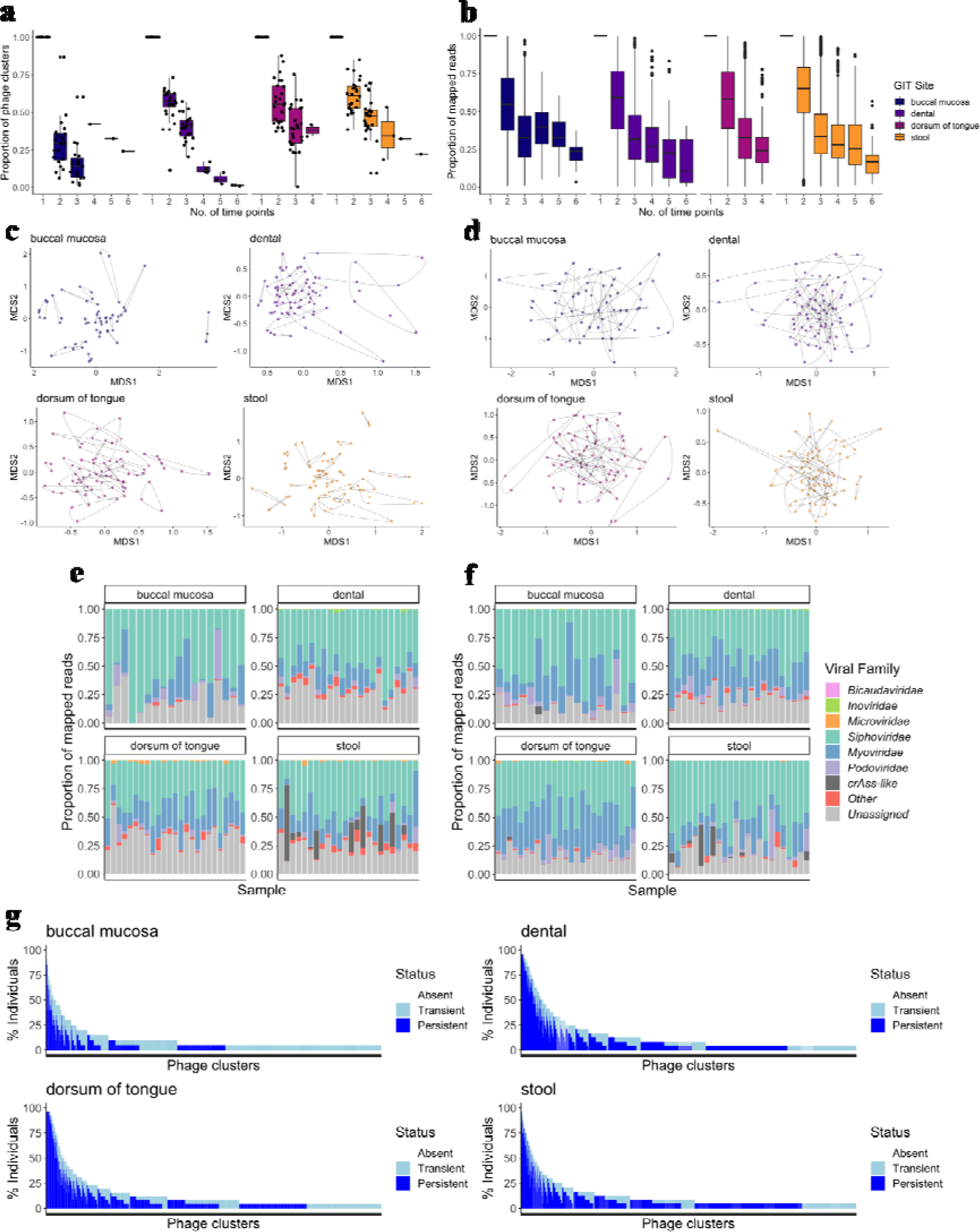
Phage cluster stability in longitudinal USA oral and gut samples. Proportion of **a)** phage clusters and **b)** reads mapped to phage clusters, in one to six timepoints for USA individuals with at least three sampling timepoints (buccal mucosa: n = 20, dental plaque: n = 24, dorsum of the tongue: n = 26, stool: n = 24). Non-metric multidimensional scaling of the Bray-Curtis dissimilarity between phage clusters incidence profiles of samples with **c)** persistent and **d)** transient phage clusters. Points represent samples and lines joining points represent grouping samples from the same individual and GIT site (buccal mucosa: n = 18, dental plaque: n = 23, dorsum of the tongue: n = 25, stool: n = 24). Proportion of reads that were mapped to phage clusters, coloured by viral family, containing only. No convergent solutions were found. Proportion of reads mapped to **e)** persistent and **f)** transient phage clusters for individuals in c) and d). “Other” represents non-phage viral families, *Alloherpesviridae, Ascoviridae, Baculoviridae, Flaviviridae, Herpesviridae, Iridoviridae, Marseilleviridae, Mimiviridae, Nudiviridae, Phycodnaviridae, Picornaviridae, Pithoviridae, Poxviridae* and *Retroviridae* (see Methods). **g)** Prevalence of transient and persistent phage clusters in the same individuals ordered by decreasing prevalence of total and persistent phage clusters.

To compare the association between stable and unstable phage clusters and GIT site, the longitudinal phageome was separated empirically into persistent and transient phage cluster profiles. Persistent phage clusters are defined as being present in three or more timepoints in a given GIT site, whereas transient phage clusters are found in less than three timepoints. There are 194 (buccal mucosa), 1377 (dental plaque), 1423 (dorsum of the tongue) and 1899 (stool) persistent phage clusters, and 1323 (buccal mucosa), 2795 (dental plaque), 3246 (dorsum of the tongue) and 2773 (stool) transient phage clusters. NMDS of persistent and transient phage cluster profiles in an ordination of two-dimensions appears to show that longitudinal samples are more clustered by individual in persistent (Fig. 3c) compared to transient profiles (Fig. 3d). PERMANOVA (permutational multivariate analysis of variance) was applied to persistent and transient phage cluster profiles for each GIT site to find which category has the highest individuality. A greater percentage of variance of persistent than transient phage clusters can be explained by individual variability across GIT sites. 78.1% (buccal mucosa), 72.3% (dental plaque), 78.4% (dorsum of the tongue) and 85.5% (stool) variance of persistent phage cluster profiles and 43.9% (buccal mucosa), 41.9% (dental plaque), 47.0% (dorsum of the tongue) and 48.6% (stool) variance of transient phage cluster profiles can be explained by individual (p < 0.001, PERMANOVA). All viral families are represented in both persistent (Fig. 3e) and transient phage clusters (Fig. 3f). Noticeably, the crAss-like family are prominently represented in persistent phage clusters of stool samples. Both persistent and transient phage clusters are found at various levels of prevalence in these individuals (Fig. 3g). The percentages of persistent phage clusters from one individual also seen in another are 76.0 ± 21.7 (buccal mucosa), 80.8 ± 5.7 (dental plaque), 81.3 ± 11.1 (dorsum of the tongue) and 75.9 ± 13.4 (stool), and for transient clusters are 69.7 ± 11.0 (buccal mucosa), 83.4 ± 7.2 (dental plaque), 81.9 ± 5.3 (dorsum of the tongue) and 76.9 ± 7.1 (stool) (median ± interquartile range). There are no significant differences in sharing of persistent or transient phage clusters between individuals for each GIT site (p = 0.477, Wilcoxon Rank Sum Test).

### Circular jumbo phages are commonly found in the oral cavity but not in the gut

Most phages have a genome size of less than 200 kbp, but we found 545 phages with genome sizes greater than 200 kbp, known as jumbo phages (Fig. 4a). 366 of these jumbo phages belong to 109 unique phage clusters, while 179 jumbo phages do not belong to any cluster (Supplementary Table 2). 97 genomes were circularised and are located in oral samples, in particular the dorsum of tongue, but not in stool (Fig. 4b). This is despite the fact that stool samples have the highest proportion of contiguous metagenomic assemblies above 200 kbp compared to oral sites (Supplementary Fig. 10). Circular jumbo phage genomes are not present in saliva from the Philippines, but this could be due to fewer assemblies (Supplementary Table 1). The largest circular jumbo phage genomes above 300 kbp are all located in dorsum of tongue samples from the USA (Fig. 4c). Of the six linear megaphage genomes above 500 kbp, one is found in a stool sample from China, and the others from three dental plaque and two from dorsum of tongue samples from the USA.

**Figure 4:**
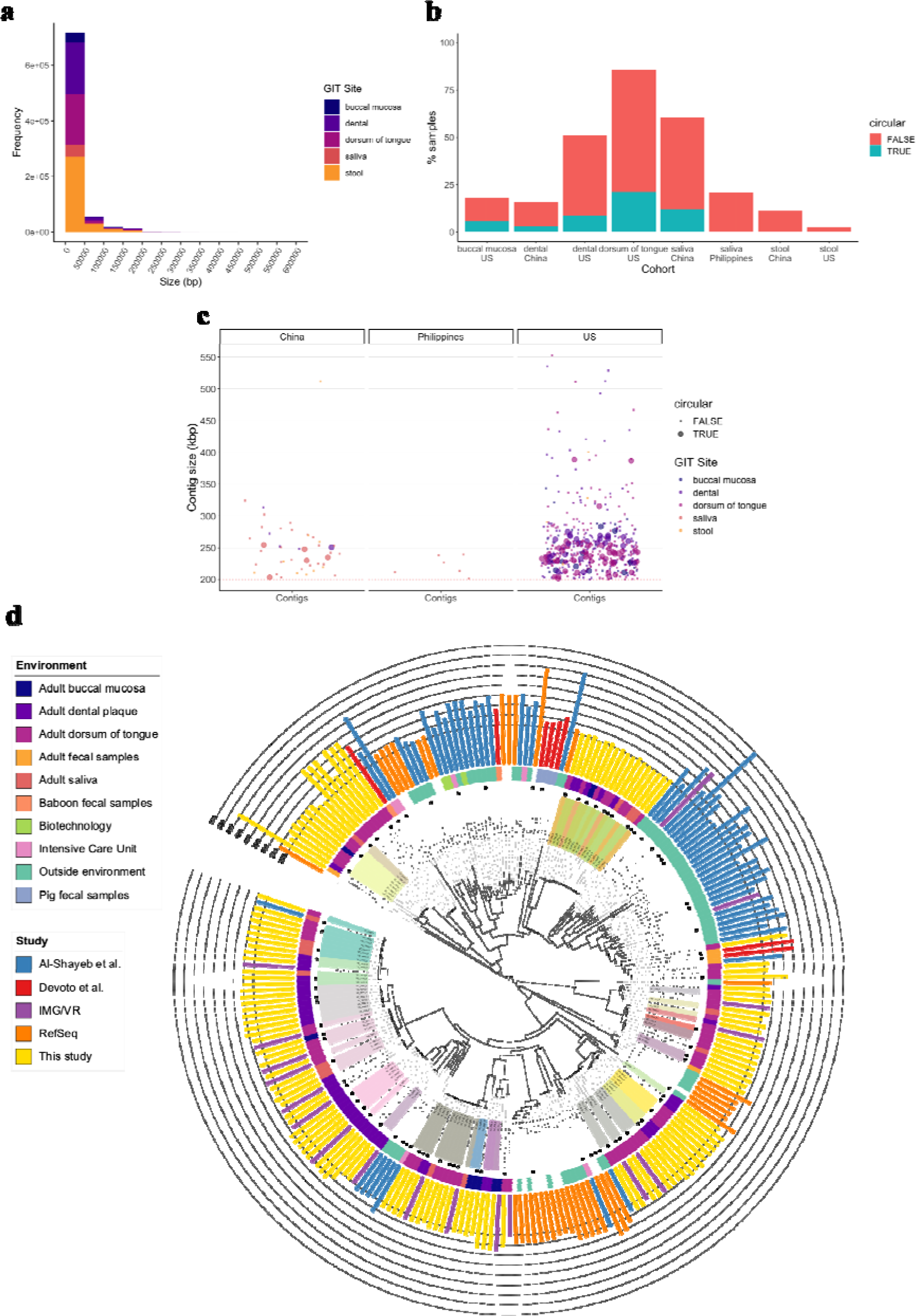
Prevalence of jumbo phages. **a)** Histogram of phage genome sizes in bp. **b)** Percentage of samples (including USA longitudinal) that contain a jumbo phage (size >= 200 kb) from USA (n = 87 buccal mucosa, n = 90 dorsum of tongue, n = 90 dental plaque and n = 70 stool), China (n = 32 dental plaque, n = 33 saliva and n = 72 stool), and the Philippines (n = 24 saliva). **c)** Sizes of unique jumbo phage contigs found in cohort. Red dashed line represents 200 kb cut-off. **d)** Phylogenetic tree of the major capsid protein (MCP) sequences of jumbo phages from this study (n = 132), RefSeq r99^19^ (n = 37), IMG/VR^56^ (n = 18), Al-Shayeb et al.^18^ (n = 56) and Devoto et al.^10^ (n = 8). Labels are coloured by phage cluster. MCPs from circularised jumbo phages that are labelled with black circles at the branch tips. The inner to outer rings show the environment and genome size with study labelled by colour. Tree bootstrapped 1000 times and rooted by midpoint.

82 circular jumbo phage genomes are found in 22 phage clusters and the remaining 15 are singletons, meaning 37 distinct circular jumbo phage groupings were identified (Supplementary Table 2). In some cases, both linear and circular jumbo phage genomes are members of the same phage clusters. 29 phage clusters containing 141 jumbo phages are persistent, i.e. found in the same GIT site in more than two timepoints. Ten phage clusters with 87 jumbo phages in the oral cavity are found across more than one country, specifically, four from China, the Philippines and the USA, five from China and the USA, and one from the Philippines and the USA. One phage cluster contains circular and linear jumbo phage genomes that are both persistent and found in all three countries.

To investigate whether the genetic variation of these jumbo phages may have arisen from other sources, the phylogeny of their major capsid proteins (MCPs) was compared with those from jumbo phages discovered in metagenomes or isolates from previous studies^10,18–20^. Large phylogenetic radiations of MCPs from oral jumbo phages intersperse with MCPs from other jumbo phages of human, animal and environmental ecosystems (Fig. 4d). Eight MCPs in jumbo phages identified from the IMG/VR database corroborate with this study’s MCPs from the same USA cohort. Notably, jumbo phages from the same clusters and gene content have the most similar MCPs. In one clade, circularised jumbo phages with related MCPs that are from different phage clusters or not belonging to a phage cluster have a conserved syntenic genome structure, but divergent amino acid alignment (Supplementary Fig. 11).

Most of the predicted protein-coding genes of these jumbo phages have hypothetical functions, while others mainly encode proteins for replication and nucleic acid metabolism (Fig. 5a, Supplementary Table 3). To investigate the possibility that these jumbo phages may carry ARGs, phage contigs were mapped against the Comprehensive Antibiotic Resistance Database (CARD)^22^. This did not reveal any jumbo phage-associated ARGs. However, auxiliary metabolic genes may provide an alternative pathway to modulating resistance to antimicrobial chemicals. Glutaredoxin, being the third most prevalent auxiliary metabolic gene, is a redox enzyme that catalyses the reduction of disulphide bonds of substrates with cofactor glutathione^23^.

**Figure 5:**
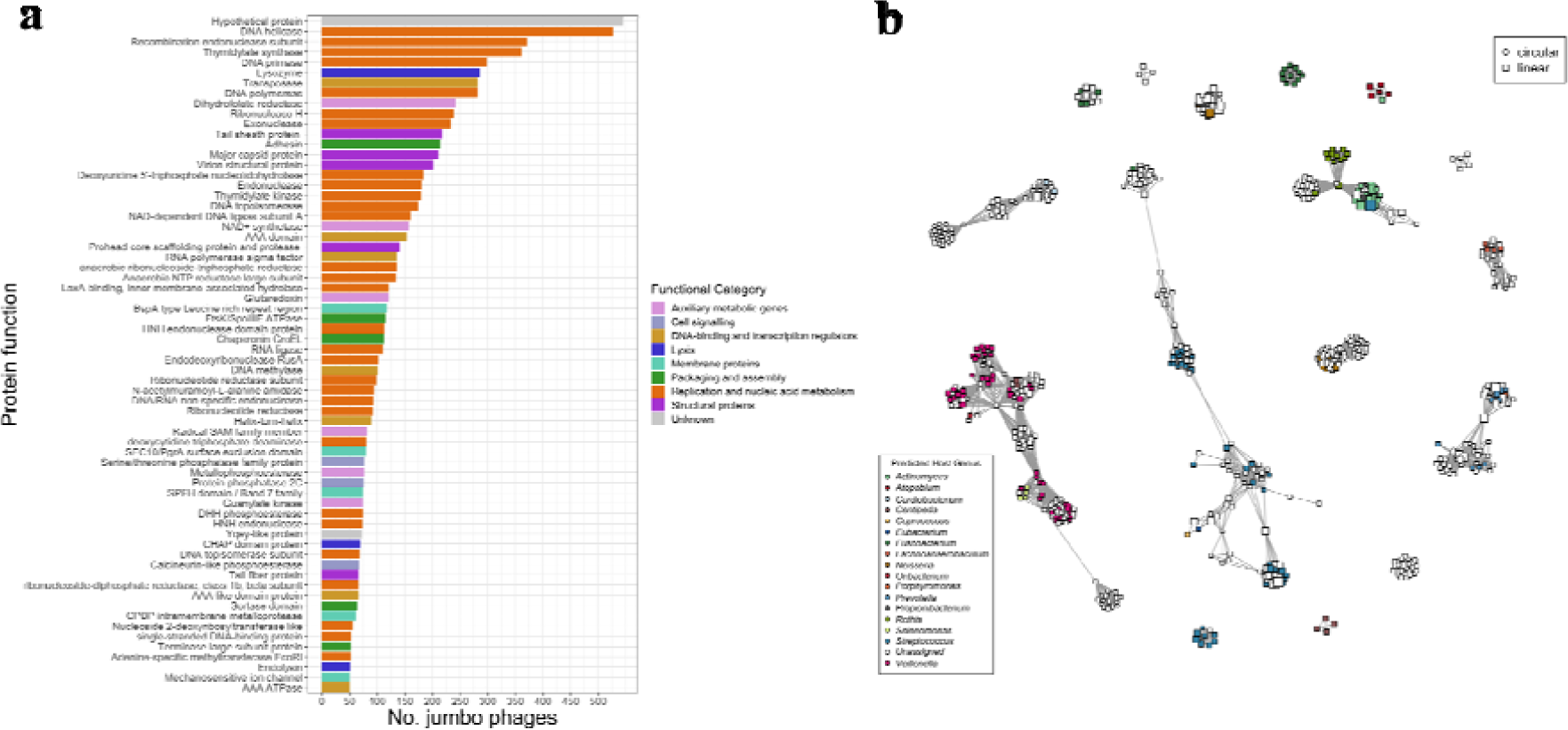
Functions and associations of protein-coding gene in jumbo phages. **a)** Number of jumbo phages (50 or greater) containing proteins coloured by functional category. Complete list of functional annotations is located in Supplementary Table 3 for each jumbo phage. **b)** Network of jumbo phages connected by ClusterONE weights of gene-sharing profiles generated from vContact2. The *Centipeda* genus represents the *C. periodontii* species^63^. Full list of jumbo phages can be found in Supplementary Table 2.

Notably, predicted host genera cluster in the same regions as jumbo phages with similar gene profiles (Fig. 5b). Using Random Forest, we identified the most important predictors among host genera by variable importance scores (Permutation Test p < 0.05). Sets of viral and bacterial proteins are important for an estimated 85.9 % (95 % CI, 77.0 – 92.3 %) accuracy of host genera (Supplementary Fig. 12). The presence of a CHAP (cysteine, histidine-dependent aminohydrolases/peptidases) domain is an important predictor of multiple hosts, particularly *Streptococcus*. CHAP domains are usually found as part of peptidoglycan hydrolases^25^. Some act as endolysins in bacteriophages that digest peptidoglycan of bacterial cell walls. It has been shown in different endolysins that CHAP domains are crucial for lytic activity^26,27^.

## Discussion

Bacteriophages are a major but largely neglected component of the oral microbiome. With their ability to introduce and transport genes between strains, as well as modulating composition through lytic activity, they have a significant impact on the population and function of the microbiome as a whole. Recent advances in bioinformatics and high-throughput sequencing now allow us to explore this community. In this study, we defined the composition of the oral phageome and compared it with the gut phageome. Firstly, we showed distinct phageome profiles between sites of the GI tract, particularly between the gut and the oral cavity, with sharing of phage clusters between proximal oral sites. Variations in phageome profiles are likely to be associated with differences in bacterial compositions of these sites that are characteristic of the gut and the oral cavity^28^. It has been observed previously that phages, particularly crAss-like, are both stable and individual-specific in the gut^16,29^. Although we did not find the same levels of crAss-like phages in the oral cavity, other phage clusters can persist for months in different oral sites and are more individual-specific than transient phage clusters. However, we cannot rule out that other less persistent phage clusters in some individuals may be attributed to lower abundance phage contigs not being picked up in our metagenomic pipeline.

Jumbo phages with circularised genomes were identified in most oral sites, particularly on the dorsum of the tongue from USA samples, but were not found in paired stool samples. The lack of circularised jumbo phages in stool samples is interesting given that stool contains large metagenomic assemblies and a higher diversity of phage clusters than saliva and buccal mucosa. Circularised jumbo phages have already been discovered in adult faecal samples from the Bangladesh, Tanzania, Peru and the USA^18,20^. However, in a recent study, 19 circularised jumbo phages were identified in saliva compared to only one in stool samples from pregnant women in the USA, suggesting the oral cavity may have a naturally higher prevalence of jumbo phages^18^.

Recombination of genetic material between bacteriophage genomes is relatively common^30^. This leads to greater variations in synteny (the physical co-localisation of protein-coding genes) particularly for larger bacteriophage genomes, meaning lineage tracing of whole genomes becomes challenging^9^. Previously, it has been shown that large terminase protein and MCP sequences of jumbo phages from environmental, human and animal samples are phylogenetically distinct from those in smaller phages of the same niches, suggesting larger phages may have arisen and evolved alongside smaller ones or that these particular sequences diverged early in evolutionary history^18^. We illustrate that MCPs are relatively conserved with gene content and synteny, which further supports the former idea that jumbo phages across ecological niches may have evolved independently.

A jumbo phage’s large genome size makes it possible for it to carry a broad range of genes^9^. We find a rich set of both viral and bacterial proteins in jumbo phage genomes of this study. No ARGs were found but instead there was a high prevalence of the auxiliary metabolic gene coding for glutaredoxin. It has been shown that *Pseudomonas aeruginosa* with mutated monothiol glutaredoxin was more susceptible to polymyxins, a last-line antibiotic for multidrug-resistant bacteria, although the exact mechanism is not known^24^. Bacteriophage genes that are homologous with bacterial genes are likely to have been acquired from host sequences during a previous infection event^31^. Lysogenic phages are also required to adapt their genetic machinery to integrate and cooperate with the host genome. This may explain why we found a relationship between the incidence of specific jumbo phage proteins and their predicted hosts. Since we could only predict bacterial hosts for 11.8 % of phages (including jumbo phages), functional protein profiles may be able to aid host predictions, especially where CRISPR spacer reference information is lacking. Phage hosts can also be refined more accurately at a metagenomic level using techniques such as SMRT metagenomic sequencing^32^, single-cell viral tagging^33^ or using *de novo* computational methods^31^.

The dorsum of the tongue contains the highest diversity of phage clusters compared to other sites, as well as the greatest prevalence of jumbo phages. This could be because bacteria in these sites are largely located in biofilms, which serve to protect microbial communities as well as phages in hostile environments, and bacteriophages themselves have been shown to promote biofilm formation^34^. Dense, protective layers of microbes and their extracellular polymeric substance matrix in biofilms could provide an opportunistic environment for the evolution of disparate phages, including jumbo phages with extended genetic machinery and metabolic capacity^9^.

Although we only find circularised jumbo phages in oral sites and a higher diversity of phage clusters on the dorsum of the tongue, we cannot rule out that the gut also may contain comparable levels of diverse phages. It is possible that the total phage composition in the gut could be misrepresented in stool metagenomes. Unlike sampling oral sites of particular physical locations, such as dental plaque, surface of the tongue and mucosa from the inner cheek, sampling faecal matter is unable to capture the spatial microbial community structure of the gut, both radially and longitudinally^35^. The microbial community structure of the human gut is of considerable importance to the dynamics of bacteriophage populations. Studies applying transmission electron microscopy of human gut biopsies have shown higher levels of bacteriophage colonisation in the colonic mucosa layer^36^ than in faeces and caecum^37^. Several existing models (piggyback-the-winner^38^ and kill-the-winner^39^) have been proposed to explain this biogeographical variation in bacteriophage density across the radial direction in the gut^40^. Lysogeny dominates in low virus to microbe ratios in the gut lumen by the piggyback-the-winner model, but switches to the lytic cycle in higher microbial densities of the mucin layer by the kill-the-winner model.

Already, significant contributions have been made in the discovery of multitudes of novel bacteriophage genomes in humans, animals, outdoor and contained environments^8,10,18,20^. However, further investigations into how genetic and functional variations are manifested at more specific biogeographical sites are required. With better structural resolution, it will be possible to capture the extent of genetic diversity and profile the dynamics of distinct phages in different human microbiomes, such as biofilms on the surface of the tongue. Clinically, there has been renewed interest in using phage therapy with other antimicrobial therapies to control biofilms^34^. In the future, combinations of genotypic attributes that influence phage persistence and interactions with their hosts could be used to select or design phages for eradicating chronic infections, like periodontitis and dental caries caused by consortia of organisms in biofilms^41^.

## Methods

### Metagenomic data for creating the phage catalogue

A total of 1061 publicly available metagenomic samples covering the USA, China and the Philippines, all sequenced using Illumina HiSeq 2000, were used to create a reference phage contig catalogue. Longitudinal USA samples were excluded from the majority of the study after the first timepoint to ensure each sample was independent, unless specified otherwise. All metagenomes passed over half the quality control metrics in FastQC 0.11.3 (https://www.bioinformatics.babraham.ac.uk/projects/fastqc/) with these pass rates calculated in MultiQC^42^. These samples include 1) longitudinal data across two years with various timepoints from the Human Microbiome Project 1 (referred to as USA) containing buccal mucosa (n = 87: 32 with one, 35 with two, 19 with three and 1 with six timepoints); dorsum of tongue (n = 91: 22 with one, 43 with two, 24 with three and 2 with four timepoints); dental plaque (n = 90: 24 with one, 41 with two, 21 with three, 1 with four and 3 with six timepoints); stool (n = 70: 13 with one, 33 with two, 21 with three, 2 with four and 1 with six timepoints)^43^, 2) healthy controls and rheumatoid arthritis (RA) patients from a Chinese study containing dental plaque (healthy: n = 32, RA: n = 76); saliva (healthy: n = 33, RA: n = 24); stool (healthy: n = 72, RA: n = 100)^44^, and 3) saliva samples (n = 24) from healthy hunter-gatherers and traditional farmers from the Philippines^45^.

Raw paired-end metagenomic reads from Chinese and Philippines samples were downloaded from the EBI (https://www.ebi.ac.uk/metagenomics/). Paired-end metagenomic samples from USA were downloaded from https://portal.hmpdacc.org/. All USA, China and Philippines samples were collected and sequenced as described in the following cited studies^43–45^.

### Processing metagenomic data

The raw reads for all samples were trimmed using AlienTrimmer v0.4.0^46^ with parameters *-k 10 -l 45 -m 5 -p 40 -q 20* and Illumina contaminant oligonucleotides (https://gitlab.pasteur.fr/aghozlan/shaman_bioblend/blob/18a17dbb44cece4a8320cce8184adb9966583aaa/alienTrimmerPF8contaminants.fasta). Human contaminant sequences were removed from all samples by discarding reads that mapped against a human reference genome (downloaded from Human Genome Resources at NCBI on 27^th^ February 2017) using Bowtie2 v2.2.3^47^ with parameters *-q -N 1 -k 1 --fr --end-to-end --phred33 --very-sensitive --no-discordant*.

### Phage contig catalogue

The reads were assembled into contigs using SPAdes v3.9.0^48^ with parameters *-k21,33,55 --only-assembler --meta*. Small contigs with a length of less than 3000 bp were removed. Linear and circular phage contigs were identified by searching for viral signatures using VirSorter v1.0.5^14^. The phage contigs were clustered with multi-alignment using BLASTn v2.6.0^49^ with parameters *-evalue 1e-20 -word_size 100 -max_target_seqs 10000* and redundant contigs were removed where identity and breadth coverage were both greater than and equal to 90%. Phage contigs were then clustered into viral clusters using vConTACT2 v0.9.10 with all default parameters^15^. Taxonomic classification of phage contigs to Order and Family level was predicted using Demovir (build 20^th^ April 2018) on phage proteins with an e-value cut-off of 1e-5 (https://github.com/feargalr/Demovir). Non-phage viral families, *Alloherpesviridae, Ascoviridae, Baculoviridae, Flaviviridae, Herpesviridae, Iridoviridae, Marseilleviridae, Mimiviridae, Nudiviridae, Phycodnaviridae, Picornaviridae, Pithoviridae, Poxviridae* and *Retroviridae*, were labelled as “Other”. Although some phage clusters were labelled as non-phage, the developers of Demovir recommend against using their classification to discriminate between bacterial, archaeal and eukaryotic sequences from metagenomic samples. The phage hosts were predicted by aligning phage contigs against a database of CRISPR spacers as described by Shkoporov et al., 2019^16^. Metadata for the phage catalogue were assembled using the helper script “create_catalogue_dataset.R”.

### Sequence and functional annotation of jumbo phages

Circular phage genomes of length > 200 kbp and linear phage genomes of length > 200 kbp that were connected to these circular phages in vConTACT2’s gene-sharing network were put forward as candidate jumbo phages using the helper script “get_candidate_jumbophages.R”. The scaffold file containing their genomes was generated using the helper script “extract_jumbophage_contigs.py”.

The candidate jumbo phage genomes were annotated for functional proteins and tRNA genes. Protein prediction was conducted using Prodigal v2.6.3^50^ with parameters *-p meta*. Protein sequences were searched against databases of Hidden Markov models, prokaryotic virus orthologous groups (pVOGs) (downloaded 1^st^ November 2019)^51^, pFAMs^52^ (downloaded 2^nd^ September 2019) and TIGRFAMs^53^ (downloaded 3^rd^ September 2019), using hmmsearch v3.2.1^54^ with e-value cut-off of 1e-5. tRNA genes were identified from nucleotide sequences using ARAGORN v1.2.36^55^ with parameters *-t -i -c -d -w* and with helper script “clean_aragorn_output.py”. The hit with the lowest e-value and domain e-value was selected from for every query protein with candidate target protein hits for each database. Next, the hit with the highest bit score and domain bit score was selected for every query protein with candidate target protein hits from more than one database. The few remaining protein query sequences with the same e-values and hit scores were deduplicated. These steps were run using the helper script “collate_functional_annotations.R”. Circular candidate jumbo phages > 200 kbp that did not contain a major capsid protein (MCP) were also excluded, leaving a total of 545 putative jumbo phages.

### Phage annotation in metagenomes

854 metagenomes from healthy individuals were mapped against the non-redundant phage catalogue using Bowtie2 v2.3.4.1. Phage contigs (excluding spurious jumbo phages) and phage clusters were quantified for each sample where contig breadth coverage was 75% or greater using helper script “phage_quantification.R”. Relative phage abundance profiles were calculated by scaling the depth coverage of phage contigs that were divided by total reads per sample. The metagenomes came from the USA with buccal mucosa (n = 87: 32 with one, 35 with two, 19 with three and 1 with six timepoints); dorsum of tongue (n = 90: 22 with one, 43 with two, 24 with three and 2 with four timepoints); dental plaque (n = 90: 24 with one, 41 with two, 21 with three, 1 with four and 3 with six timepoints); and stool samples (n = 70: 13 with one, 33 with two, 21 with three, 2 with four and 1 with six timepoints), China with dental plaque (n = 32); saliva (n = 33); and stool samples (n = 72), and the Philippines with saliva samples (n = 24). Metadata for the samples can be found in Supplementary Table 1. The following analysis was conducted in script “phage_analysis.R”.

### Phage diversity

To find differences in beta-diversity of phage profiles between groups of individuals, the Bray-Curtis dissimilarity of phage incidence (presence or absence) profiles was computed between individuals and visualised using NMDS. Silhouette analysis of k-medoids was used to select the number of distinct groups with the largest Silhouette width.

The alpha-diversity was calculated as the phage cluster richness which is the number of unique phage clusters for each sample. Only samples with greater than 100 phages were included. Individuals with samples containing less than or equal to three clusters were excluded from the group comparison. The phage cluster richness was compared between paired GIT sites from the same individuals in each of the following groups: China dental plaque vs. saliva (n = 30); China stool vs. saliva (n = 30); China stool vs. dental plaque (n = 30); USA buccal mucosa vs. dental plaque (n = 45); USA buccal mucosa vs. dorsum of tongue (n = 45); USA buccal mucosa vs. stool (n = 36); USA dental plaque vs. dorsum of tongue (n = 86); USA dental plaque vs. stool (n = 67); and USA dorsum of tongue vs. stool (n = 68). Since the number of phage contigs in each sample is significantly linearly correlated with Phage Cluster Richness (p < 2.2×10^−16^) (Supplementary Fig. 13), the number of phage clusters for each sample were subsampled to the smallest number of phages found in a sample for each paired comparison. The phage cluster richness between samples in each group was tested for statistical significance with a Two-sided Wilcoxon Rank Sum Test.

### Microbial composition

MetaPhlAn2 v2.6.0^17^ was used to identify the composition of bacteria and archaea from samples apart from longitudinal USA samples. One dorsum of tongue USA sample did not have bacterial nor archaeal microbial predictions. Procrustes analysis was applied to visualise the superposition of NMDS dimensions of phage incidence profiles on microbial genera incidence profiles using the protest function in the vegan package v2.5.6 in R. PROTEST was performed with 999 permutations to a significance of p = 0.001.

### Longitudinal analysis of phages

The stability of phages was investigated by computing the number of timepoints each phage cluster is found from each individual and GIT site in the longitudinal USA data. The proportion of phage clusters and reads mapped against these clusters were calculated for each number of sampled timepoints available for each individual and GIT site: buccal mucosa (n = 20), dental plaque (n = 24), dorsum of the tongue (n = 26) and stool (n = 24). Persistent phage clusters are defined as being found in three or more timepoints, whereas transient phage clusters are defined as being found in less than three timepoints. The Bray-Curtis dissimilarity was computed between persistent and transient phage cluster incidence profiles from GIT sites of individuals containing both persistent and transient phage clusters: buccal mucosa (n = 18), dental plaque (n = 23), dorsum of the tongue (n = 25) and stool (n = 24). Non-metric multidimensional scaling was applied to scale the dissimilarity into a two-dimensional ordination using the metaMDS function in the vegan package v2.5.6 in R. PERMANOVA analysis was performed using the adonis function in the vegan package.

### Antimicrobial resistance gene annotation of jumbo phages

Phage contigs were annotated for ARGs by mapping against CARD v3.0.0 using BLASTn v2.10.0 with parameters *-evalue 1e-5*. Hits were filtered by 90% identity.

### Phylogenetic analysis of MCPs

A phylogenetic tree was constructed from MCP sequences of jumbo phages identified from this study, complete dsDNA jumbo phage genomes from RefSeq r99^19^, and jumbo phage scaffolds from IMG/VR^56^, Al-Shayeb et al.^18^ and Devoto et al.^10^. MCPs from IMG/VR jumbo phage scaffolds were identified using the functional annotation pipeline described above. Jumbo phages from Al-Shayeb et al. that contain MCPs were identified using the helper script “get_files_with_MCPs.sh” from supplementary Genbank files, generated as described in Al-Shayeb et al.^18^. MCPs of complete jumbo phages were also collected from Genbank files downloaded from RefSeq r99. All MCPs were collated into a file of amino acid sequences using the helper script “get_MCPs.py”. The MCP amino acid sequences were clustered using CD-HIT^57^ at 100% identity to remove redundancy. The output sequences were aligned using MAFFT v7.453^58^ using parameters *--localpair --maxiterate 1000*. The phylogenetic tree was constructed using IQTREE v1.6.12^59^ with automatic model selection and 1000 bootstrap replicates, and visualised using iTOL v5.5.1^60^ with the tree rooted at midpoint. Metadata of these jumbo phages for the tree was created using helper scripts “get_RefSeq_info.py”, “create_tree_metadata.R” and “add_metadata_to_tree_annotations.sh”.

### Synteny comparison

Pairwise alignments of genomes of four circular jumbo phages with closely related MCPs (SRS078425_NODE_1_length_226348_cov_18.355, SRS019607_NODE_9_length_235695_cov_219.599, ERR589420_NODE_6_length_229992_cov_13.7426, SRS044662_NODE_25_length_232905_cov_15.7753) were performed using tblastx v2.10.0 within Easyfig v2.2.2^61^. Genbank files of the circular jumbo phages were generated using RASTtk v1.3.0^62^ and separate genbank files were created for the four jumbo phages using the helper script “extract_genbank_files.py”. MCPs with type “CDS” in their Genbank files were relabelled “MCP”. Genomes were linearised and fixed by MCP as the starting protein-coding gene using the helper script “reorder_genbank_files.py”.

### Phage Host Prediction

A Random Forest classification model was created to analyse the relationship between functional profiles in jumbo phages and their host genera. Incidence of protein function for 109 phage clusters containing 366 jumbo phages and 179 jumbo phages with no assigned phage cluster, that have CRISPR spacer-predicted hosts, were used to train the model using the randomForest package v4.6-14 in R. The accuracy of the trained model was evaluated using the in-built out-of-bag error estimation. The Mean Decrease in Accuracy was determined to measure the importance of protein functions for predicting host genera. Its significance was estimated by permuting the host genera response variable to produce null distributions of the Mean Decrease in Accuracy (scaled by the standard error) for each protein function predictor using the rfPermute package v2.1.81.

### Code availability

The code for the analysis is available from https://github.com/APC-Microbiome-Ireland/phageome_analysis

## Acknowledgements

The research was supported by the Centre for Host-Microbiome Interactions, King’s College London, funded by the Biotechnology and Biological Sciences Research Council (BBSRC) grant BB/M009513/1 awarded to D.L.M., The Alan Turing Institute under the Engineering and Physical Sciences Research Council (EPSRC) grant EP/N510129/1, and APC Microbiome Ireland funded by the Research Centre grant from Science Foundation Ireland (SFI) under Grant Number SFI/12/RC/2273. The authors would like to thank Professor Gordon B. Proctor, Dr Saeed Shoaie and Dr Sunjae Lee for constructive criticism of the manuscript.

## Author contributions

V.R.C. and C.H. conceived the presented idea. A.S. created the phage catalogue. A.S. and V.R.C. did the data analysis. V.R.C. and A.S. wrote the manuscript with support from C.H., D.L.M., D.G-C and P.M.

