## Supplementary Material for "The human oral phageome is highly diverse and rich in jumbo phages"

**Supplementary Figures**

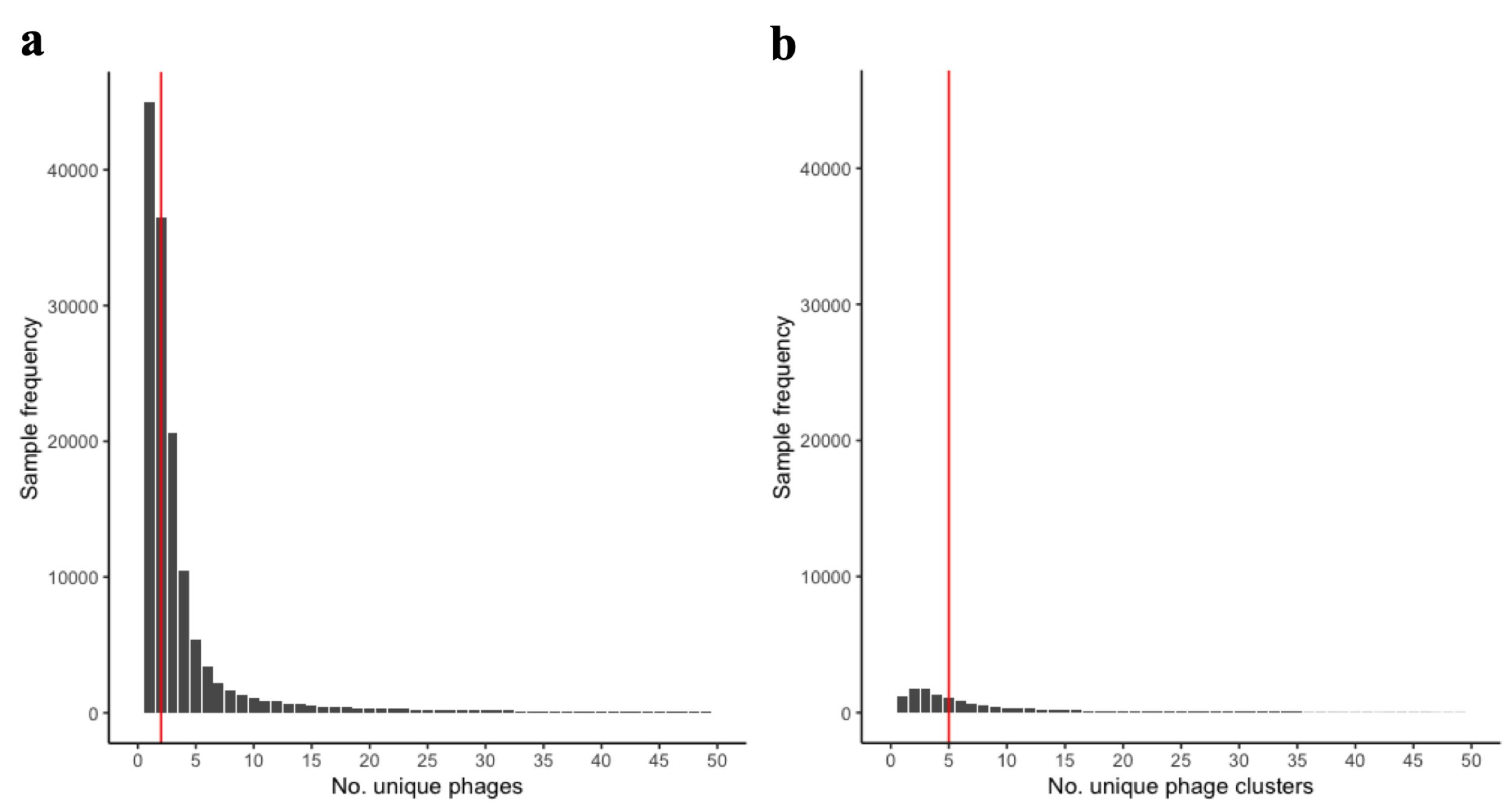

**Supplementary Figure 1:** Frequency of samples containing a number of unique a) phages and b) phage clusters. Red line represents median number.

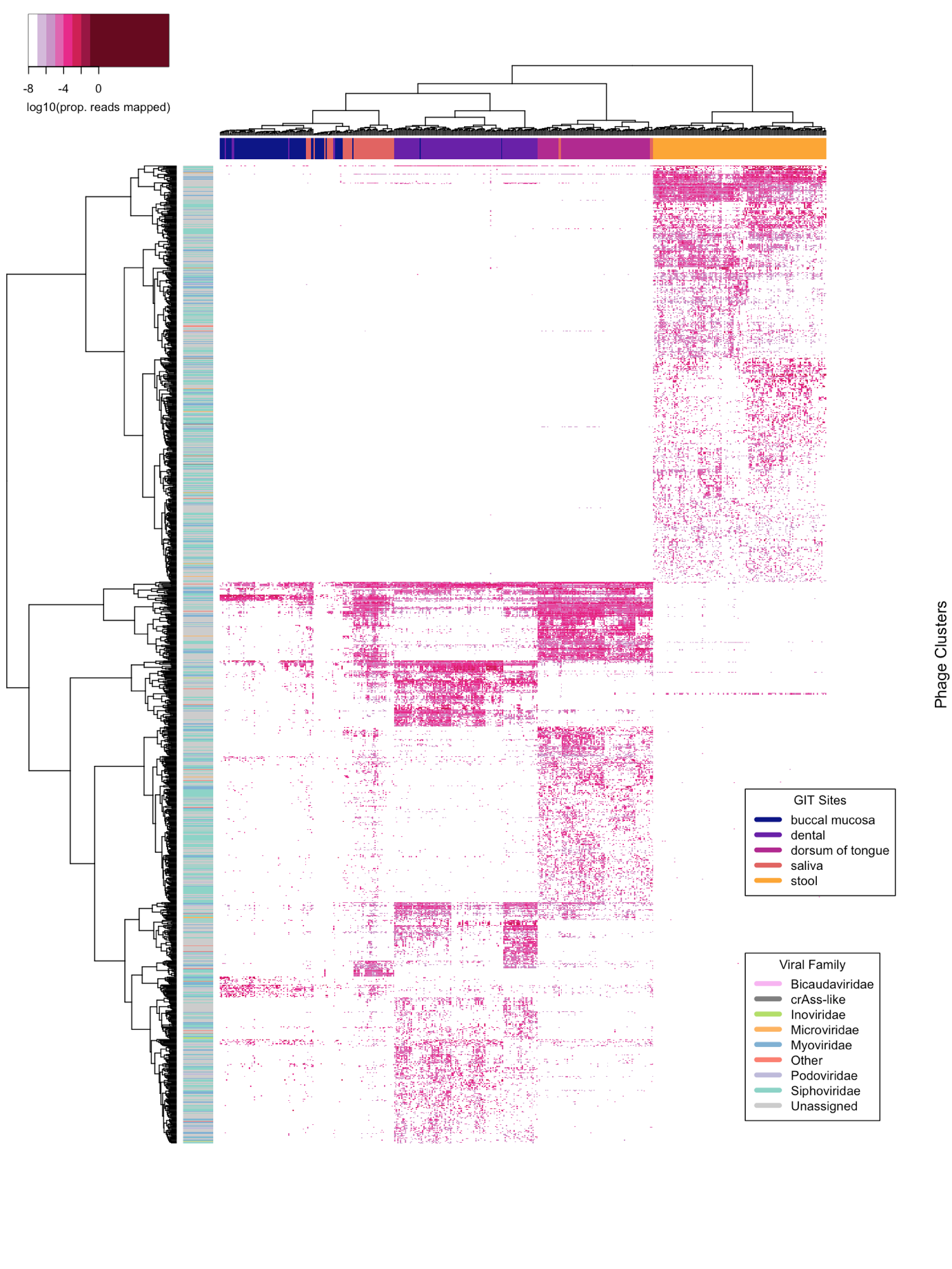

**Supplementary Figure 2: Log10 of the proportion of reads mapped to differentially abundant phage clusters for each sample**. Samples and phage clusters clustered by hierarchical clustering. Phage clusters that were differentially abundant between groups were selected where Bonferroni-corrected adjusted p-values < 0.001 from Kruskal-Wallis Rank Sum test. USA buccal mucosa (n = 87), dorsum of tongue (n = 90), dental plaque (n = 90) and stool (n = 70); China dental plaque (n = 32), saliva (n = 33) and stool (n = 72); and Philippines saliva (n = 24).

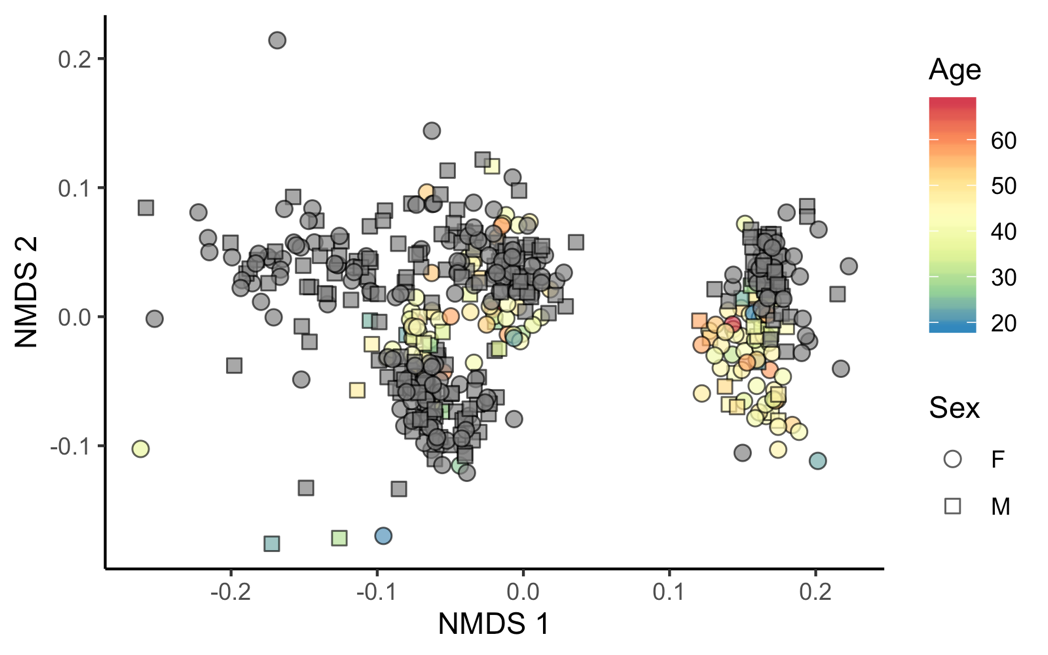

**Supplementary Figure 3: Phage incidence and abundance profiles.** Non-metric multidimensional scaling of Bray-Curtis dissimilarity between phage cluster incidence profiles of samples (excluding longitudinal USA) labelled by GIT site and geographical location.

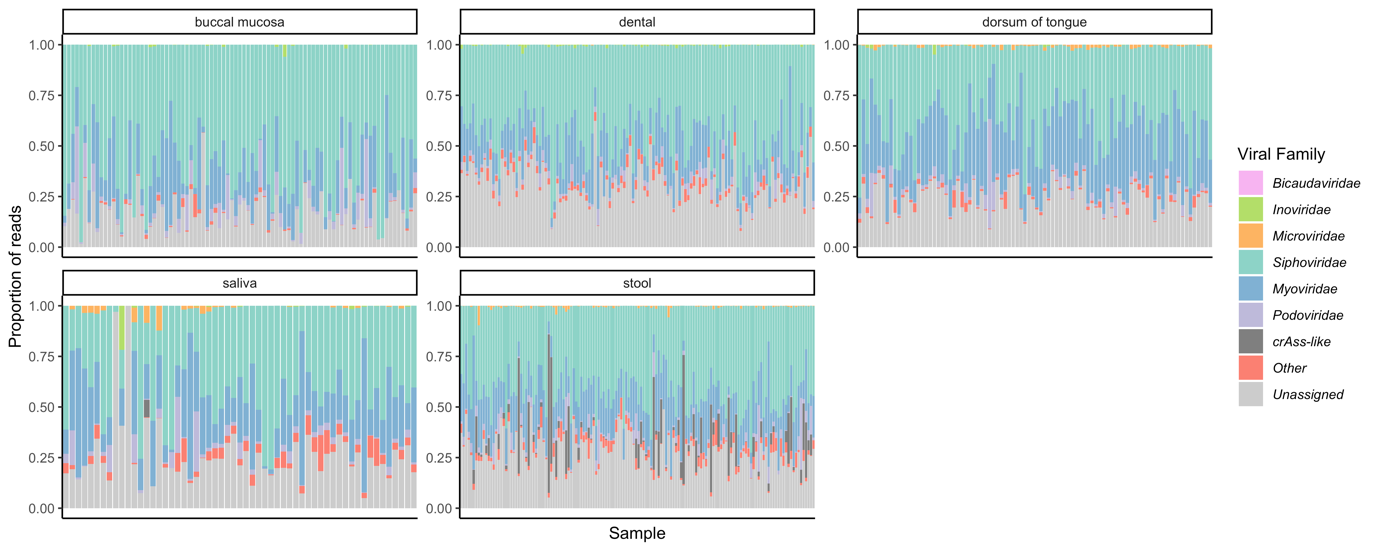

**Supplementary Figure 4: Relative abundance of phage taxonomy across GIT sites.** Proportion of total reads that are mapped to phage clusters, coloured by viral family, for each sample (excluding USA longitudinal). “Other” represents non-phage viral families, *Alloherpesviridae*, *Ascoviridae*, *Baculoviridae*, *Flaviviridae*, *Herpesviridae*, *Iridoviridae*, *Marseilleviridae*, *Mimiviridae*, *Nudiviridae*, *Phycodnaviridae*, *Picornaviridae*, *Pithoviridae*, *Poxviridae* and *Retroviridae* (see Methods). USA buccal mucosa (n = 87), dorsum of tongue (n = 90), dental plaque (n = 90) and stool (n = 70); China dental plaque (n = 32), saliva (n = 33) and stool (n = 72); and Philippines saliva (n = 24).

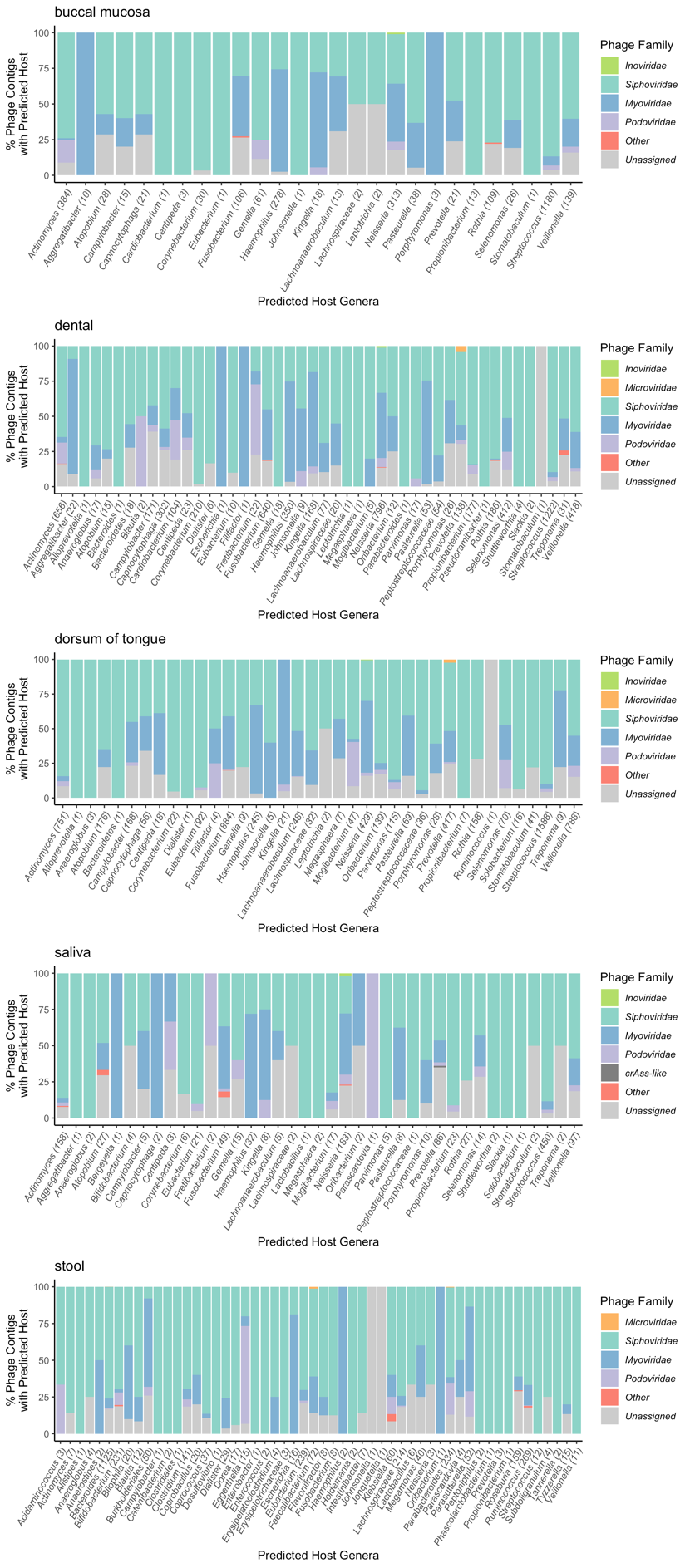

**Supplementary Figure 5: Percentage of phage contigs of phage families with predicted bacteria hosts.** 16,513 (out of 139,929) phage contigs were assigned to host genera. Bracketed number after genus name represent the total number of phage contigs assigned to a particular genus.

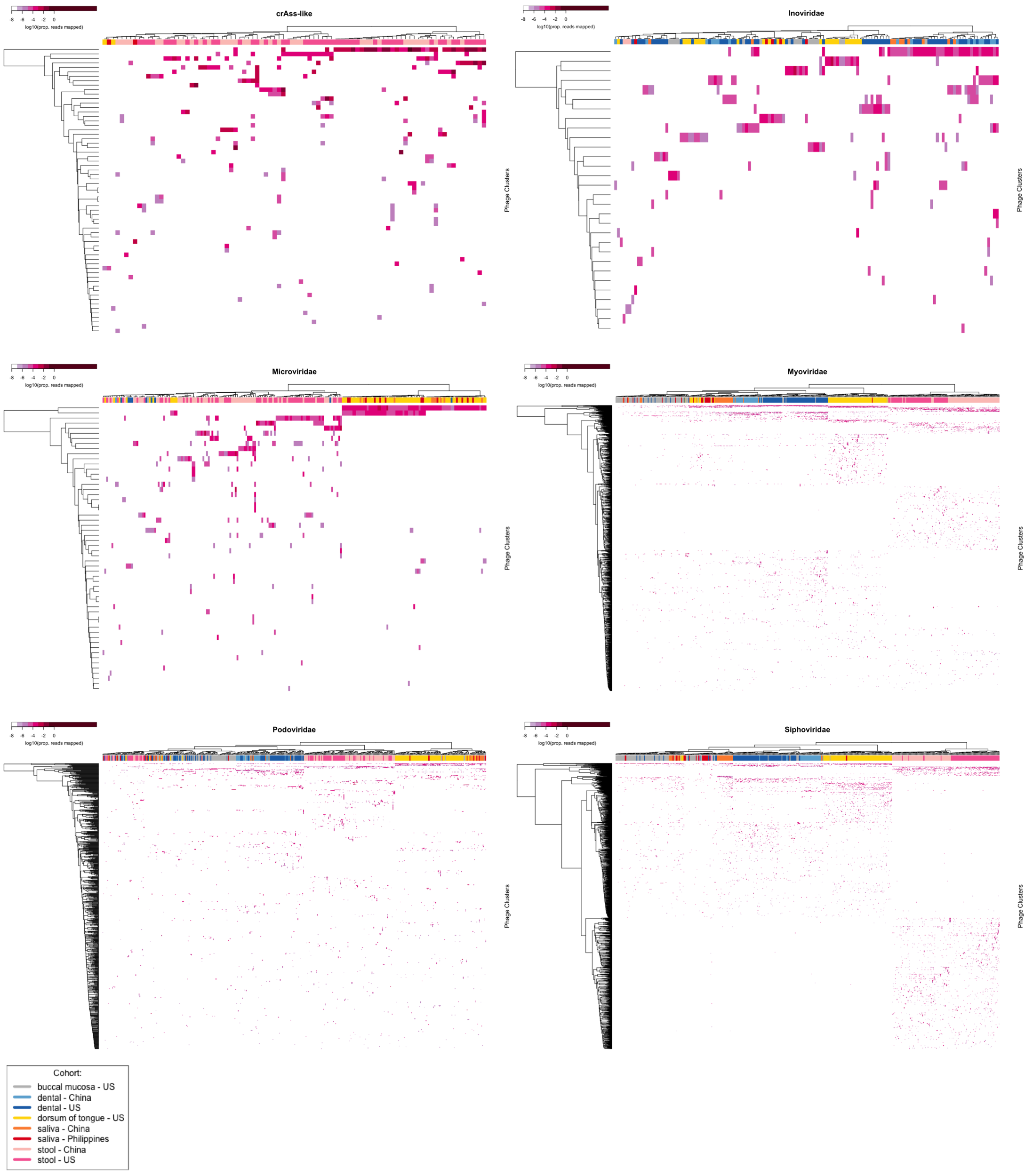

**Supplementary Figure 6: Log10 relative abundance of phage clusters for *crAss-like*, *Inoviridae*, *Microviridae*, *Myoviridae*, *Podoviridae* and *Siphoviridae* phage families.** Hierarchical clustering of all samples (excluding USA longitudinal). *Bicaudaviridae* was only found in USA buccal mucosa (n = 87), dorsum of tongue (n = 90), dental plaque (n = 90) and stool (n = 70); China dental plaque (n = 32), saliva (n = 33) and stool (n = 72); and Philippines saliva (n = 24).

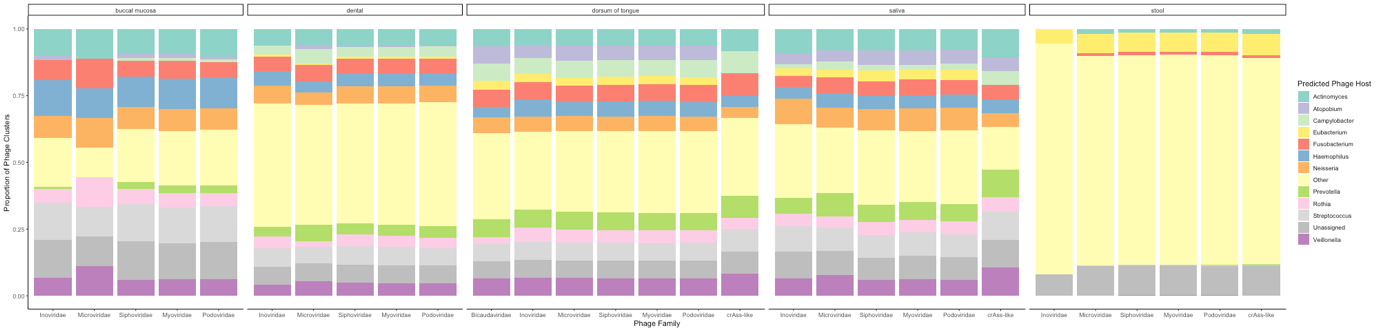

**Supplementary Figure 7: Proportion of phage clusters with predicted phage hosts for each phage family and GIT site.** Buccal mucosa (n = 87 from the USA), dorsum of tongue (n = 90 from the USA), dental plaque (n = 90 from the USA and n = 32 from China), saliva (n = 33 from China and n = 24 from the Philippines) and stool (n = 70 from the USA and n = 72 from China) (Excluding longitudinal USA samples).

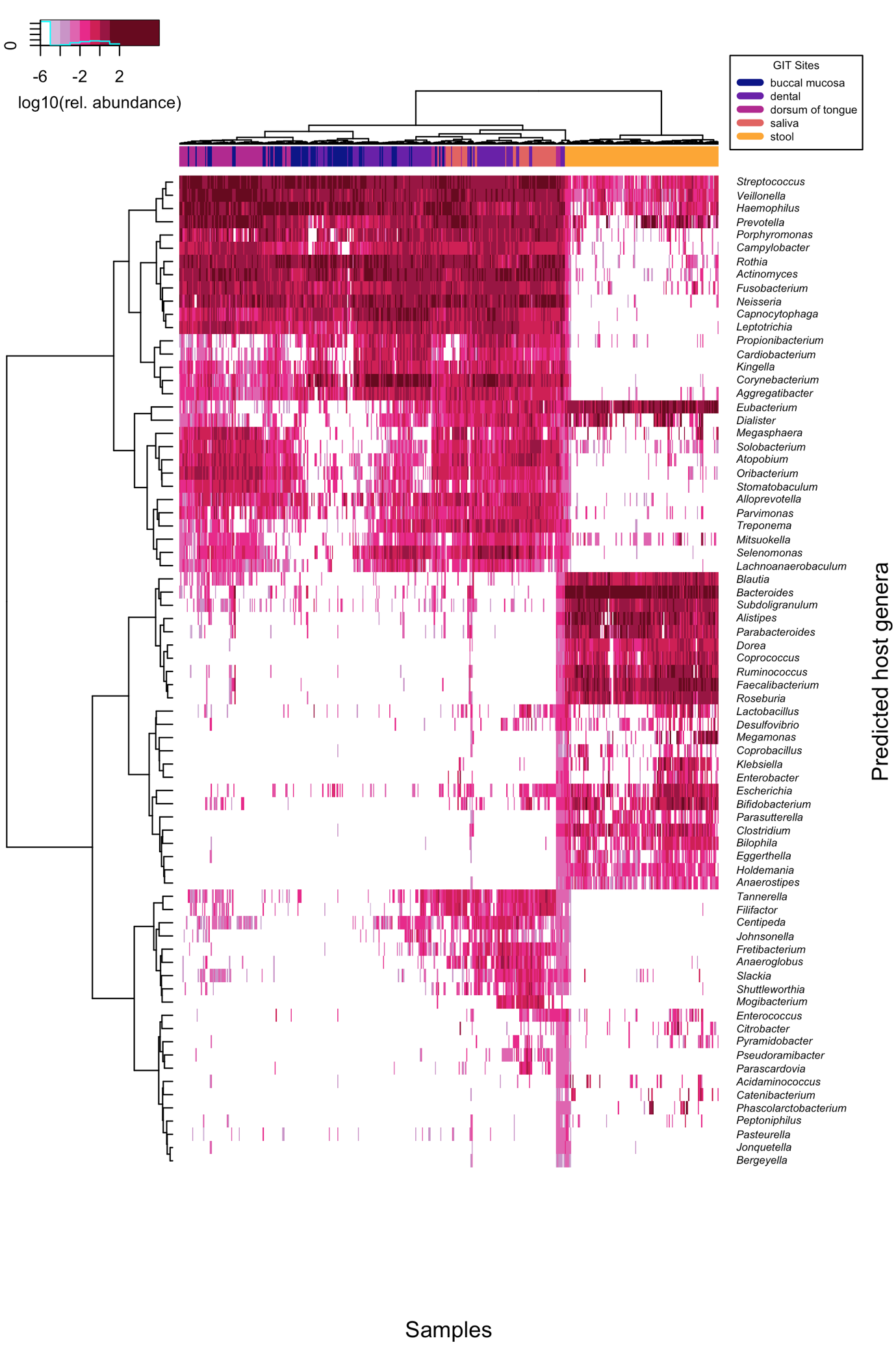

**Supplementary Figure 8: Log10 relative abundance of bacterial genera.** Hierarchical clustering of buccal mucosa (n = 87 from the USA), dorsum of tongue (n = 90 from the USA), dental plaque (n = 90 from the USA and n = 32 from China), saliva (n = 33 from China and n = 24 from the Philippines) and stool (n = 70 from the USA and n = 72 from China). (Excluding longitudinal USA samples).

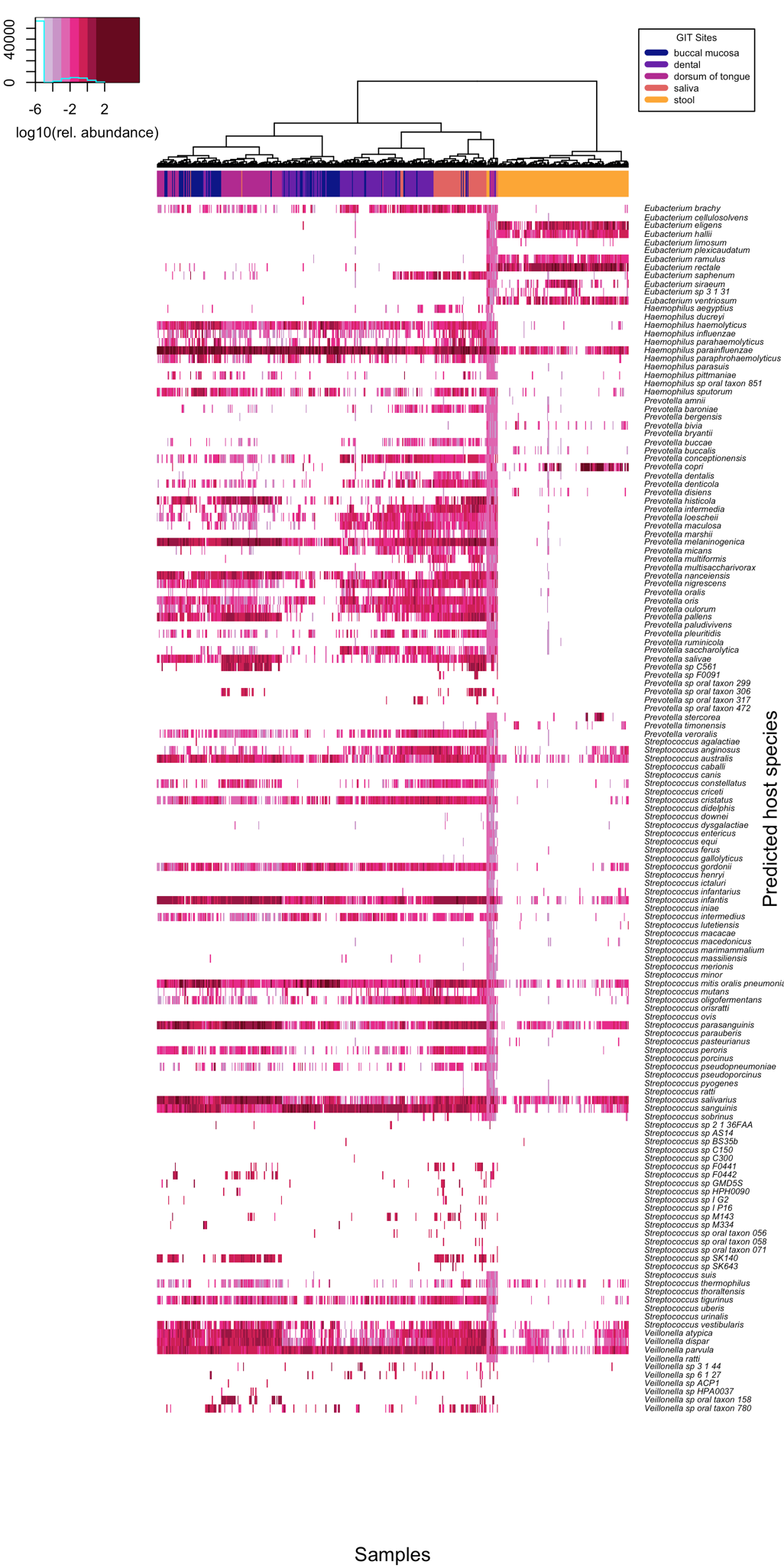

**Supplementary Figure 9: Log10 relative abundance of *Eubacterium*, *Haemophilus,* *Prevotella*, *Streptococcus* and *Veillonella* species.** Hierarchical clustering of buccal mucosa (n = 87 from the USA), dorsum of tongue (n = 90 from the USA), dental plaque (n = 90 from the USA and n = 32 from China), saliva (n = 33 from China and n = 24 from the Philippines) and stool (n = 70 from the USA and n = 72 from China). (Excluding longitudinal USA samples).

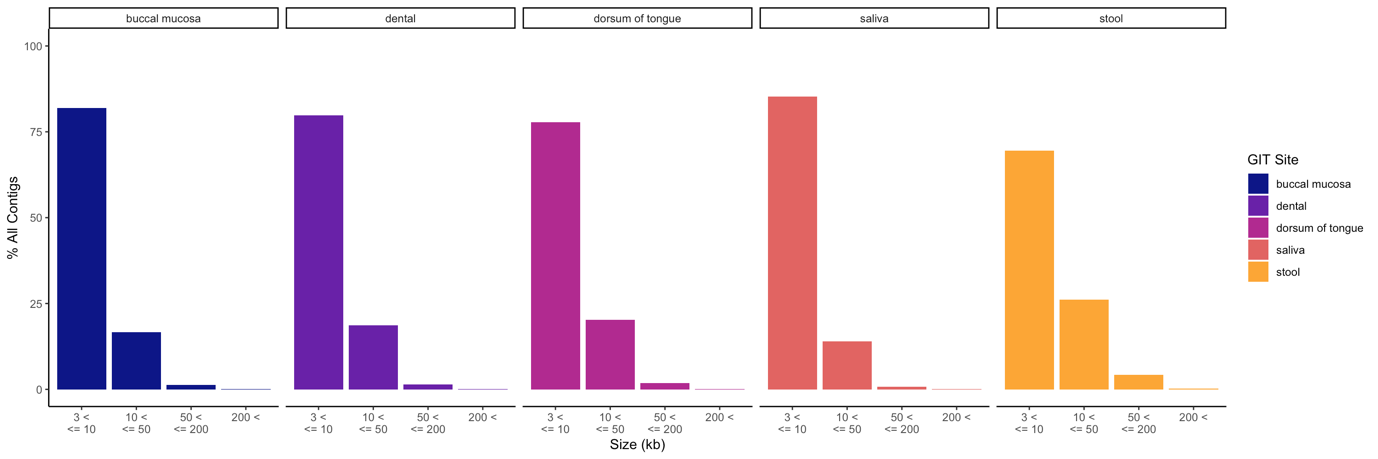

**Supplementary Figure 10: Contig size.** Percentage of all contigs (phage and non-phage) that have genome sizes between 3 – 10, 10 – 50, 50 – 200 and over 200 kb for each GIT site

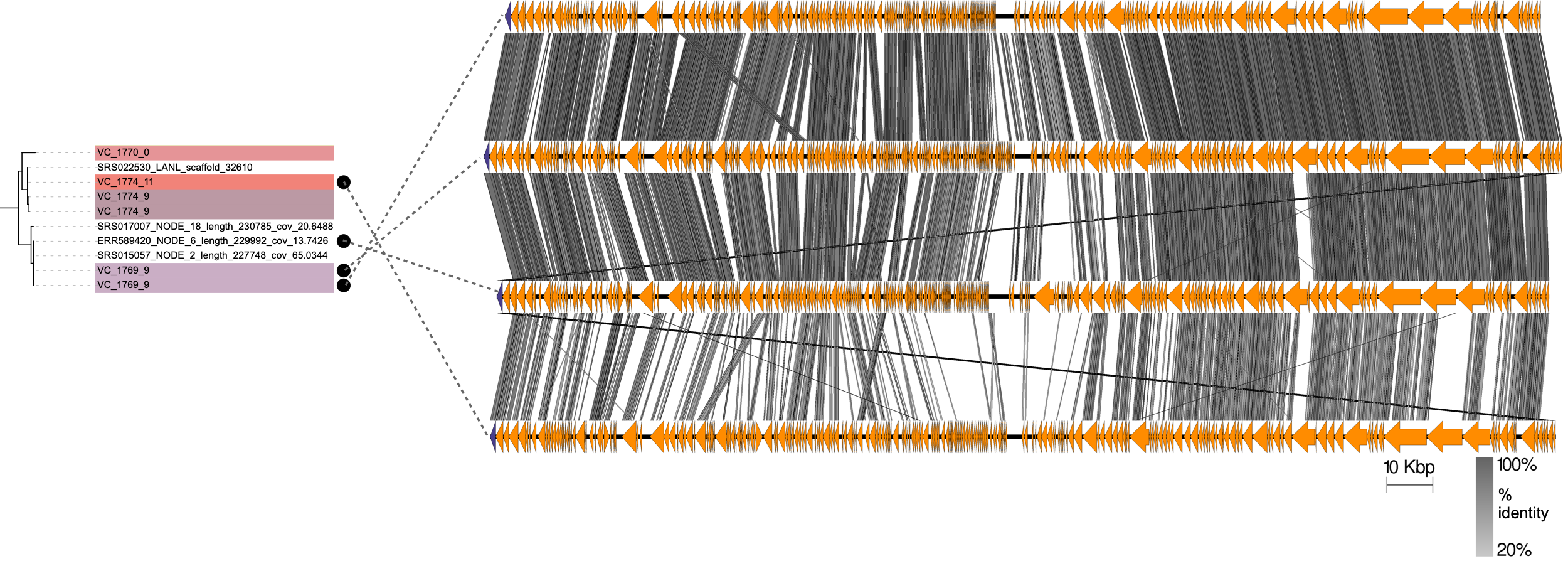

**Supplementary Figure 11: Comparison of genomic loci of circular jumbo phages from a clade of related MCPs**. Circular genomes are linearised and fixed by their MCPs (shown as purple arrows). Other protein-coding genes are shown as orange arrows. Alignment was performed using tblastx (See Methods).

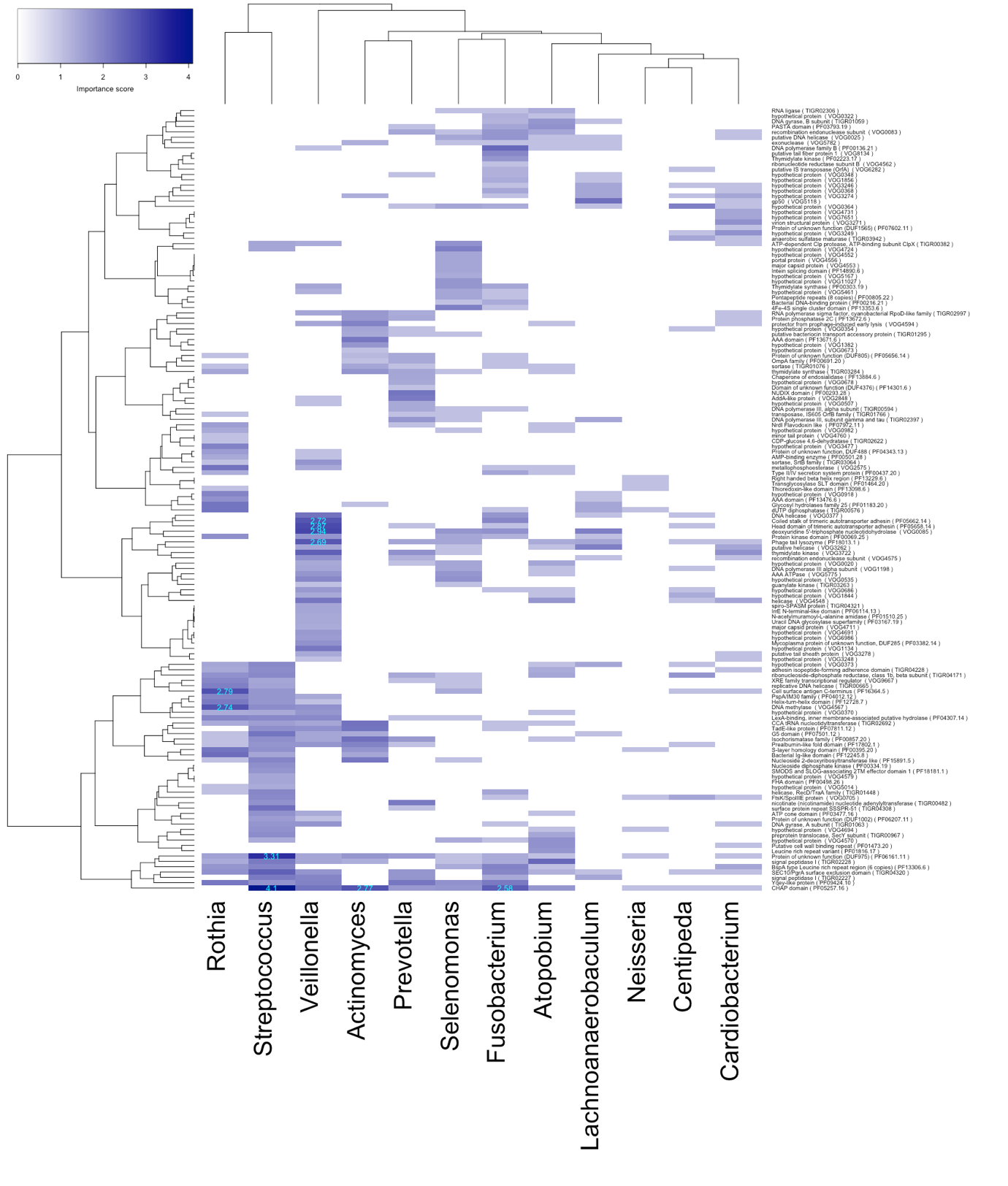

**Supplementary Figure 12:** Importance score measured as the Mean Decrease in Accuracy (scaled by S.E.) for each protein and host genus from a Random Forest model where Permutation Test p-value < 0.05 for each individual and all out-of-bag cross validated predictions (See Methods). Rows labelled by protein description and pVOG/pFAMs/TIGRFAMs identifiers. The highest ten importance scores are highlighted in light blue

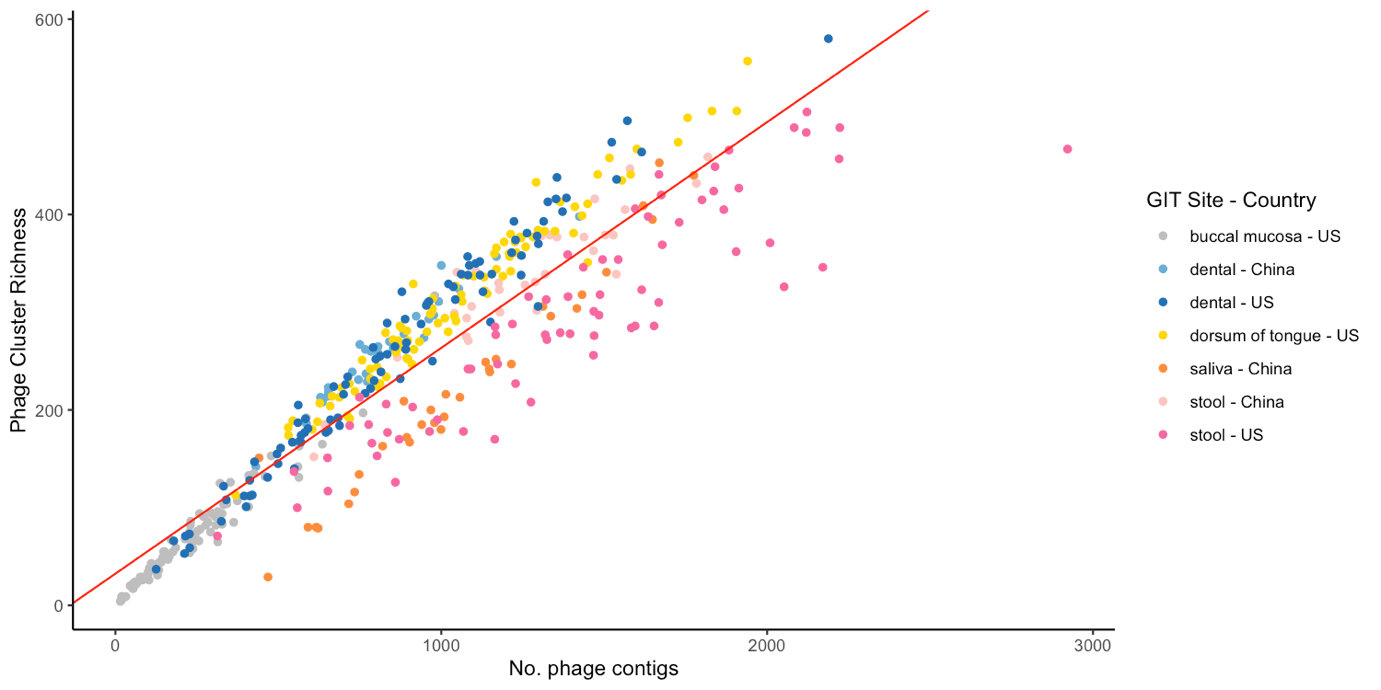

**Supplementary Figure 13:** Linear regression of Phage Cluster Richness against number of phage contigs in each sample from China and the USA (Adjusted R^2^ = 0.8618, p-value < 2.2x10^-16^)
